# Tuning cell motility via cell tension with a mechanochemical cell migration model

**DOI:** 10.1101/847046

**Authors:** K. Tao, J. Wang, X. Kuang, W. Wang, F. Liu, L. Zhang

## Abstract

Cell migration is orchestrated by a complicated mechanochemical system. However, few cell migration models take account of the coupling between a biochemical network and mechanical factors. Here, we construct a mechanochemical cell migration model to study the cell tension effect on cell migration. Our model incorporates the interactions between Rac-GTP, Rac-GDP, F-actin, myosin, and cell tension, and it is based on phase field approach hence very convenient in describing the cell shape change. This model captures common features of cell polarization, cell shape change, and cell migration modes. It shows cell tension inhibits migration ability monotonically when cells are applied with persistent external stimuli. On the other hand, if random internal noise is significant, the regulation of cell tension exerts a non-monotonic effect on cell migration. As the elevation of cell tension impedes the formation of multiple protrusions hence enhances the streamline position of the cell body. Therefore the migration ability could be maximized at intermediate cell tension under random internal noise. These model predictions are consistent with our singlecell experiments and other experimental results.

**Statement of significance:** Cell migration plays a vital role in many biological processes such as tumor metastasis. It is a complicated process regulated by dynamic coupling between the biochemical network and mechanical forces. However, few cell migration models take account of both factors. Here, we construct a mechanochemical cell migration model to study how cell migration is regulated by cell tension. Our model predicts that cell tension not only inhibits cell movement under persistent external stimuli but also prompts cell migration under random internal noise when cell tension is low. Therefore an optimized cell tension could maximize the migration ability under random internal noise. We further confirmed these model predictions are consistent with our single-cell experiments and other published experimental results.

## Introduction

Cell migration plays a vital role in many biological processes, including tumor metastasis, embryonic morphogenesis, tissue repair, and regeneration. In most eukaryotic cells, one of the salient features in terms of migration is the combined effect from actin filament polymerization against cell membrane and the action of molecular motors to retract the rear of the cell. The polymerization of F-actin results in the reorganization of the cytoskeleton, which controls the formation of the lamellipodium. This process of cytoskeletal rearrangement is regulated by multiple chemical factors, notably Rho family GTPases (1). The effective Rho GTPase positively enhances the activity of cytoskeleton effector proteins through integrated signaling pathways, resulting in a myriad of processes such as actomyosin movement (2). Another factor taken into account is cell membrane tension, which provides global feedback to inhibit the propulsion of actin filaments (3–7). Moreover, membrane bending rigidity is also relevant especially for cell collisions with an obstacle or other cells.

Cell migration is regulated by dynamic coupling between the biochemical network and mechanical forces. Cytoskeletal rearrangements and the mechanical forces, i. e., cell tension, are not only downstream outputs of biochemical cascades but also exerts feedbacks on biochemical signaling networks (8, 9) to enable cells to make cellular decisions. The cross-talk between biochemical network and mechanical forces exists over different spatial scales (10). At the sub-cellular scale, both the signaling events (11, 12) and the patterned activity of actomyosin lead to protrusive or contractile dynamics of the cortical cytoskeleton, which generate forces that ultimately integrate whole-cell behaviors like cell shape changes and migrations (13). While at cellular scale polymerized F-actin contributes to part of cell membrane tension and the increase of tension reversely stall actin assembly by mechanical means (14, 15).

However, despite numerous mathematical models on cell migration, few of them take account of coupling between biochemical signals and mechanical factors. Many one-dimensional models were established (16–18) to capture specific aspects of cell migration, e.g., thickness and flow profile of lamellipodium (16). For two-dimensional models, different models are applied for different cell types. Lee *et. al*. introduced Graded Radial Extension (GRE) to explain the steady motion of fish keratocytes (19). This model considers a sharp interface of cell membrane, which assumes the extension of the front and retraction of the rear of keratocytes occurring perpendicularly to the cell edge (20). Another approach developed for modeling the chemotactic behavior of social amoeba and human neutrophils was the level set method, which has the advantage of describing featureless interfaces, without tension or other physical properties. For diffusive interface models, the popular one was phase field model that can couple actin flows with physical forces and was applied into a wide range of cell types to investigate migration mechanisms (21–23). Phase field model has also been applied to study multiple cell motion behaviors such as membrane fusion (24) and cell delamination (25). Although detailed mechanisms for the cytoskeletal mechanics are unknown, many reviews have covered this aspect. Lim et. al. provide a summary of continuum-based models of the mechanical stiffness of cells (26). There are also reviews for cell mechanics, such as cytoskeleton (27–29), actin protrusion (30), or cell signaling and cell migration (31).

In our previous work (3), we established a mechanochemical model on cell polarization and performed single-cell experiments to show that the elevation of tension inhibits the polarization tendency and enhances the persistence of polarity. Cell polarization precedes cell migration. Here, we extend this mechanochemical model to study cell migration via tunable cell tension. Our cell migration model incorporates the interactions between Rac-GTP, Rac-GDP, F-actin, myosin and cell tension, and it is based on phase field model hence very convenient in describing cell shape change. We demonstrate that our mechanochemical cell migration model captures two typical cell morphological patterns, yields three migration modes with high to low migration capability under random internal noise, and persistent movement mode under external stimuli. And when cell tension is lower, cell shape deformation is more significant and cell migration capability is higher. Our model predicts that cell tension inhibits protrusion hence stalls cell movement under persistent external stimuli. It also shows that elevated cell tension could prompt cell migration as directional migration can be impeded by the formation of multiple protrusions under random internal noise when cell tension is low. Therefore migration ability could maximize at intermediate cell tension under random internal noise. We further confirm that these model predictions are consistent with our singlecell experiments and other experimental results.

## Materials and methods

### Mechanochemical migration model

A cell is considered as a round disk Ω_0_ with the radius *R* = 5 *μm*, which is of the actual size and imbedded into a larger 2D computational domain Ω with a size of 40 *μm* × 40*μm*. The choices of all parameters are listed in the S1 Table. The diffusion coefficients of *D_u_* and *D_v_* are set according to experimental results. Some parameters (i.e., the basal conversion rate *b*, maximum activation rate *c*_3_, dephosphorylation rate *r*, disassociation constants *K_i_*(*i* = 1,2,3), etc.) are based on the published papers. We use parameter fitting to determine the values of all the other parameters, including the maximum activation/inhibition rate *c_i_*(*i* = 1,2,4,5) and equilibrium constants in Hill function *K_i_*(*i* = 4,5). The degradation rates of Factin and myosin are estimated via our previous work.

Our model includes protein interactions between four species, namely polarity marker protein in active form Rac-GTP (*u*) and inactive Rac-GDP (*v*), polymerized Factin (*f*) and motor protein myosin (*m*). Notably, the wave pinning (WP) model provides a minimal reaction-diffusion system with bi-stable kinetics to pin the waves into a stable polar distribution on expressing protein interactions in a modeling framework (32). Therefore, we extend the WP model to describe interchange between Rac-GTP on the cell membrane and Rac-GDP is in cytosol as well as their interaction with F-actin and myosin in cytosol. Since molecules move more freely in the cytoplasm than on membrane, the ordering between diffusion coefficients *D_u_*, *D_v_*,*D_m_* and *D_f_* is *D_u_* < *D_f_* ≈ *D_m_* ≪ *D_v_*. The basal conversion rate from Rac-GDP to Rac-GTP is denoted as *b* and inversely Rac-GTP dephosphorylates to Rac-GDP is with the rate of *r*. The total molecule number of Rac-GTP and Rac-GDP is conserved. Self-activation of Rac-GTP and the positive feedback from F-actin are described by non-linear Hill functions 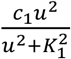 and 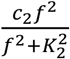 with *c*_1_ and *c*_2_ representing the maximum self-activation rate while *K*_1_ and *K*_2_ meaning microscopic disassociation constants. Likewise, the mutual inhibition of F-actin and myosin are expressed by the form of 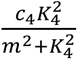 and 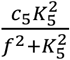 with *c*_4_ and *c*_5_ being the maximum self-production rate of Factin and maximum inhibition rate respectively, while *K*_4_ and *K*_5_ denoting equilibrium constants. Meanwhile, degradation rates of F-actin and myosin are denoted as *d_f_* and *d_m_* respectively.

In our two-dimensional (2D) model, cell tension (denoted as *mt*) is regarded as a global inhibition factor that comprises of cortical tension part and bilayer membrane part. In a unit length of membrane, cortical tension is proportional to the total amount of polymerized F-actin, while membrane tension is determined by the cell shape. Hence the total tension can be derived as *mt*(*f*) = *σ*(1 + ∫_Ω_0__*fdxdy*) *L*, where *σ* represents initial line membrane tension and *L* represents cell perimeter. F-actin polymerization is activated by Rac-GTP as 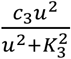, where *c*_3_ is the maximum regulation rate and again *K*_3_ is the disassociation constant. Furthermore, this process is also down-regulated by cell tension, i.e., proportional to 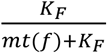, where *K_F_* is a scaling factor for non-dimensionalization.

Based on the descriptions above, the dynamics of the model system is expressed by the following equations.

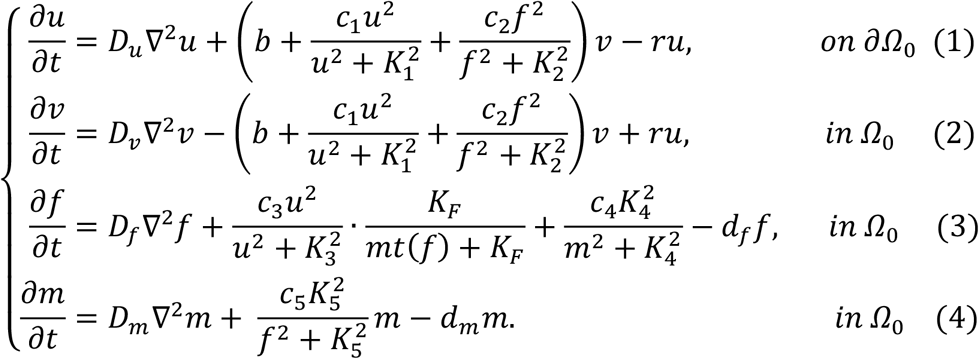

Considering Ω_0_ constantly changes its shape due to deformation from various forces, we use phase field model to track cell interface by introducing an order parameter ϕ to distinguish Rac-GTP (*u*) on *∂Ω*_0_ from Rac-GDP (*v*), F-actin (*f*), and myosin (*m*) in Ω_0_.

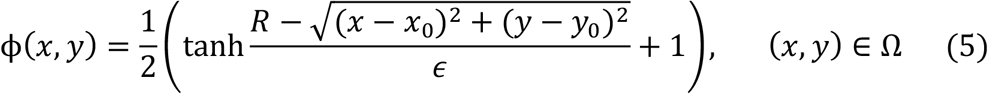

where (*x*_0_, *y*_0_) is the initial position of the cell. Thus two phases (ϕ(*x,y*) = 1, and ϕ(*x,y*) = 0) are naturally distinguished via this order parameter. The cell membrane is described by a narrow transition layer between the interior of the cell (ϕ(*x,y*) = 1) and the exterior of a cell (ϕ(*x,y*) = 0) with a width of *ϵ*. The position of the cell membrane is approximated by *B*(ϕ) = 3ϕ^2^(1 − *ϕ*)^2^, which vanishes outside the narrow interface. By coupling the order parameter, Eqs. 1–4 can be re-arranged as follow.

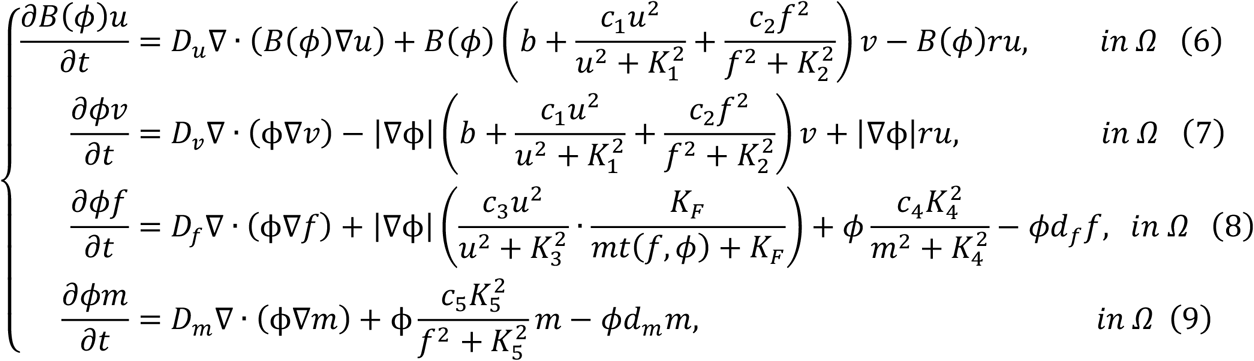

where 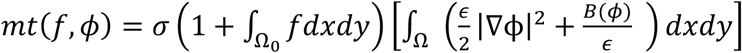 since the cell perimeter *L* is approximated as 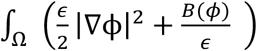 in the diffusive interface.

The shape of cell membrane is determined by interactions of various forces, such as the tension force from cell tension *mt*(*f, ϕ*), which converts from area density to line density (22) and yields

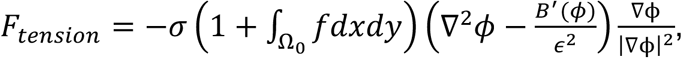

and the pressure from volume conservation constraints which is defined as

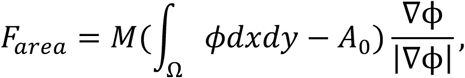

where *M* is the penalty coefficient and *A*_0_ is the prescribed area. The movement of the cell membrane is due to protrusion force from polymerized F-actin and contraction force from myosin

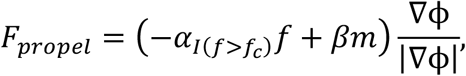

where *α* and *β* are protrusive and contractive coefficient respectively. Note that *I*(·) is an indicator function assuming protrusion exists only if the concentration of F-actin is beyond a critical value *f_c_*, since actin filaments form part of cytoskeleton first and then the excessive polymerized actin induces protrusion. In simulations, we presume *f_c_* = 0.8*f_max_*, where *f_max_* represents the maximum concentration of F-actin under the steady state. Also, adhesiveness from the attachment or detachment from the substrate can be viewed as friction, which is defined as *F_friction_* = −*τV*, where *τ* is friction coefficient and *V* is the velocity of membrane motility on a normal direction. Meanwhile, we assume the motion of the cell membrane is overdamped and thus satisfies force balance criteria, meaning *F_tension_* + *F_area_* + *F_propel_* + *F_friction_* = 0. Since the evolution of phase field ϕ follows *∂*_t_ϕ = −*V* · *∇ϕ*, we obtain the final equation

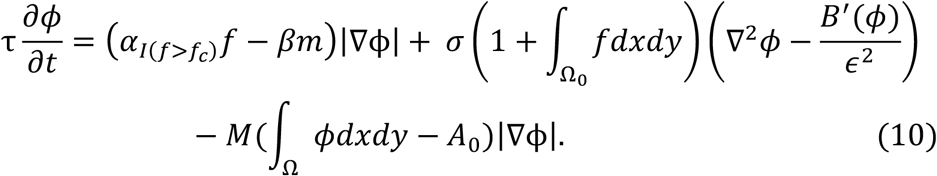

Eq. 10 expresses that the inertial force friction, which is transmitted from the substrate to cells is balanced by active forces and the movement of membrane results in cell motility. Note that we neglect the momentum transfer from F-actin to the substrate and the area constraint as the perimeter of cell is not conserved from experiments.

The external stimulus is applied to Eq. 6 and Eq. 7 to accelerate the rate of transformation from Rac-GDP to Rac-GTP. In simulations, the stimulus is expressed as *k_s_v* and *k_s_* are spatially dependent. The setup of an external stimulus is similar as discussed in our previous work.

The global graded stimulus *k_s_* is

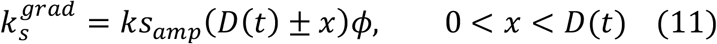

where *ks_amp_* means the amplitude of the stimulus and *D*(*t*) is the maximum radius of cells which depends on time *t* because cell changes morphology and moves during simulations. In the beginning, *D*(0) = 2*R*, which is the diameter of cells at the initial shape. We also define the local stimulus as

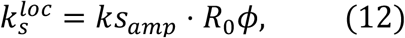

where *R*_0_ is a uniformly distributed random number between 0 and 1. Both global graded stimulus and local stimulus are persistent in simulations.

We choose the periodic boundary condition and apply the semi-implicit Fourier-spectral method for the spatial discretization to solve Eq. 6 to Eq. 10. The initial conditions affect the spatial profile of Rac-GTP. If the initial concentrations of Rac-GTP and Rac-GDP are too low, their distributions are unlikely to break symmetry and therefore the cell remains non-polarized regardless of the strength of the stimulus, let alone cell deformation and motility. Homogeneous initial conditions of *u* = 2*μm*^−2^, *v* = 6*μm*^−2^, *f* = 0, *m* = 2*μm*^−2^ are provided.

### Cell line and cell culture

The human breast cancer cell lines, MCF-7 and MDA-MB-231, were purchased from American Type Culture Collection (ATCC). MDA-MB-231 cells were stably transfected with the plasmids coding for 3-phosphoinositide-specific Akt-PH domain fused with CFP and the plasmids coding for Rac1 fused with YFP using FuGENE^®^ HD transfection reagents (Promega). MCF-7 cells and MDA-MB-231 cells were cultured in Dulbecco’s Modified Eagle’s Medium (DMEM) supplemented with 10% fetal bovine serum (FBS), 100 *μg/mL* penicillin, and 100 *μg/mL* streptomycin at 37°C in a 5% CO_2_ atmosphere mixed with 95% air. MCF-7 cells were stained with anti-CD44-PE (BD Pharmingen), and anti-CD24-Alexa Fluor 488 (BioLegend) antibodies for sorting with a flow cytometer (BD FACS Aria II). The CD44^+^/CD24^-^ phenotype was regarded as CSCs, whose proportion was approximately 1.6% (33); the other three phenotypes, CD44^+^/CD24^+^, CD44^-^/CD24^+^, and CD44^-^/CD24^-^, represented the NSCCs (34, 35). CSCs sorted from MCF-7 cells were cultured in DMEM/F12 (1:1 mixture of DMEM and Ham’s F-12 medium) with 20 *ng/mL* basic fibroblast growth factor (bFGF), 10 *ng/mL* epidermal growth factor (EGF), B27 serum-free supplement and N-2 supplement to inhibit the differentiation of CSCs (33). The phenotype of CSCs was maintained for at least one month.

### Imaging and image processing

Before loading cells, the culture dish was incubated with Geltrex (120-180 μg/mL, Gibco) at 37°C for an hour. MCF-7 cells cultured in the 24-well plate were stained with 1 *μg/mL* Hoechst 33342 for 20 minutes at 37°C and rinsed with PBS. Subsequently, they were stained with 333 *μg/mL* Golgi-Tracker Red (Molecular Probes) at 4°C for 30 minutes, rinsed with ice-cold medium and incubated in fresh medium for 30 minutes at 37°C. Then, the cells were loaded into confocal culture dishes to measure the movements of spontaneous polarization. Time-lapse imaging was performed on a Zeiss inverted microscope (Axio Observer.Z1 (SP)) every 5 minutes for at least 1 hour. The objective was 20X with an NA=0.5 in air. The detection channels included bright-field, DAPI, and mCherry using an X-cite lamp.

MDA-MB-231 cells were harvested by trypsinization and then seeded in the 6-well glass bottom cell culture plates coated with Geltrex at the density of 1700 *cells/cm^2^* in DMEM supplemented with 1% FBS medium (isotonic medium). The samples were incubated at 37°C for 12 hours. The medium was replaced by fresh isotonic medium, hypotonic medium (a 1:1 mixture of isotonic medium and double distilled water), which increased cell tension (36), or medium contained deoxycholate (Sigma) at a concentration of 400 *μM*, which decreased cell tension (36). After 2 hours, the plates were mounted in the motorized stage of a Nikon Eclipse Ti-E wide-field fluorescence microscope to record cell deformation and migration. The objective was 20X with an NA=0.75 in air, and the intermediate magnification was 1.5X. The detection channels included bright-field, YFP, and CFP. Thirty regions of interest in each well were selected and images were recorded every 6 minutes for least 2 hours. Then fresh isotonic medium, water or deoxycholate was added into the plates to adjust the cell tension. After 2 hours, thirty new regions of interest in each well were selected and images were recorded every 6 minutes for another 2 hours. Quantifications of cell deformation, velocity and motion direction were carried out using the Matlab.

## Results

### The mechanochemical cell migration model captures common features of cell polarization

We confirm that the new cell migration model maintains the common features of cell polarization as our previous cell polarization model. First, the simulations show F-actin filaments congregate at one side of the polarized cell at a steady state in response to a gradient external stimulus. And the maximum concentration of F-actin decreases from 0.7 *μm*^−1^ to around 0.3 *μm*^−1^ as cell tension increases from 0.2 *pN*/*μm* to 0.8 *pN*/*μm* (Fig. 1 b). Second, cells polarize only if the amplitude or duration of the external stimulus is above a certain threshold. And these thresholds increase at higher cell tension, e.g., the threshold of the duration at fixed amplitudes triples when cell tension is doubly elevated (Fig. 1 c). Third, cells with lower tension have a higher tendency to polarize (Fig. 1 d), consistent with the experimental results that membrane tension serves as a global inhibitor of cell polarization,

**Fig. 1.**
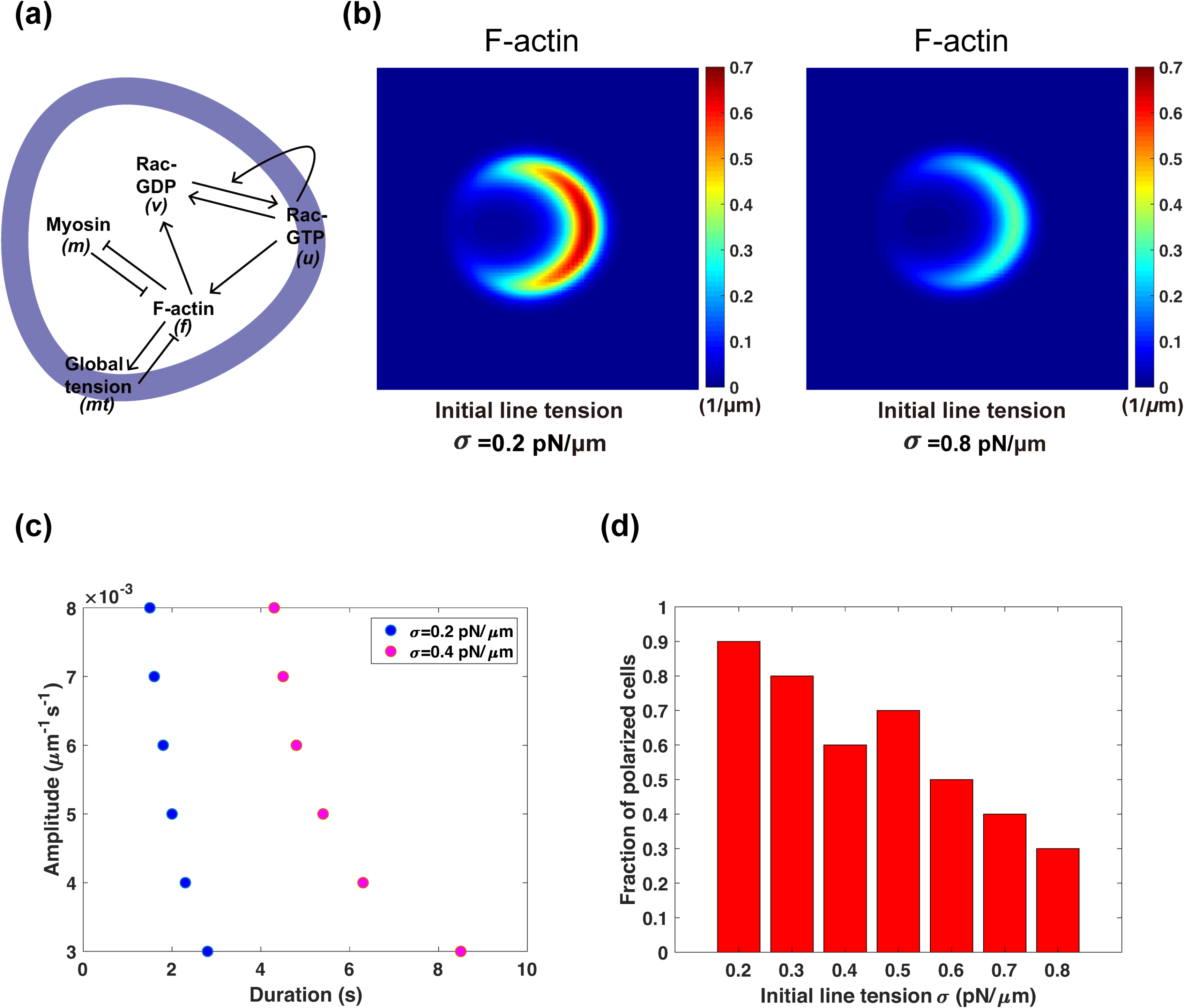
Characterization of the cell polarization occurring ahead of migration in the mechanochemical cell migration model. **(a)** Schematic diagram of the regulatory network in the cell migration model. Cell tension (*mt*) globally inhibits (denoted by—|) the formation of F-actin (*f*) in the cell. However, F-actin polymerization increases (black solid arrow →) cortical part of cell tension, and meanwhile activates membrane-bound Rac-GTP (*u*) (light blue). Rac-GTP is also activated by stimulation and itself. In addition, Rac-GTP on the membrane and Rac-GDP (*v*) in the cytosol interconvert. Myosin(*m*) attached to the actin bundles is mutually antagonized with F-actin. **(b)** The steady-state spatial profiles of F-actin in a polarized cell with initial line tension *σ* = 0.2 *pN*/*μm* and 0.8 *pN*/*μm* respectively. To respond to a gradient external stimulus, F-actin accumulates at the cell front in the cytoplasm. As *σ* rises to 0.8 *pN*/*μm*, the maximum concentration of F-actin decreases. **(c)** The threshold of the amplitude of stimuli as a function of the duration of stimuli for inducing cell polarity at different values of initial line tension. **(d)** The fraction of polarized cells under various values of *σ*. Each bar counts the results from repetitive simulations under the sample size *N* = 20.

### The deformation of the cell membrane under different cell tension

Polarized cells are capable of protruding outward and form lamellipodiums toward the directions where F-actin has polarized distributions. As a result, cell membrane is deformed. Just like cell polarization, the deformation of cell membrane should be strongly influenced by cell tension. The maximum concentration of F-actin is significantly low in cells with larger tension (Fig. 1 b) and the probability of cell polarization drops at higher cell tension.

Our cell migration model demonstrates cell membrane deformation given the random internal noise, i.e, the varying conversion rate from Rac-GDP to Rac-GTP, same as the local stimulus defined in Eq. 12 with *R*_0_ to be an *N*(0,1) Gaussian distribution random variable. And the model displays two typical types of morphological patterns when the initial line tension is attuned as 0.005 *pN*/*μm* and 0.015 *pN*/*μm* accordingly (Fig. 2 a and c, Movie S1 and S2). Initially, the cell is round with an even distribution of F-actin. In one type, F-actin starts to form several clusters because of the positive feedback from Rac-GTP and cytoskeleton is slightly changed (Fig. 2 a, *t* = 20 *min*). Eventually, F-actin clusters disperse in a larger area with the maximum concentration increases and the cell membrane has distinct protrusions (Fig. 2 a, *t =* 60 *min)*. Depending on the random stimuli, cells present various morphologies. In the other type of patterns, F-actin stays at low concentrations and is distributed more homogeneously, meaning the cell is unpolarized (Fig. 2 c). And cells keep a stable shape. Moreover, the tendency of the two types of patterns varies at different cell tension. The first one has more tendency to be observed at low cell tension since the cell membrane is more flexible.

**Fig. 2.**
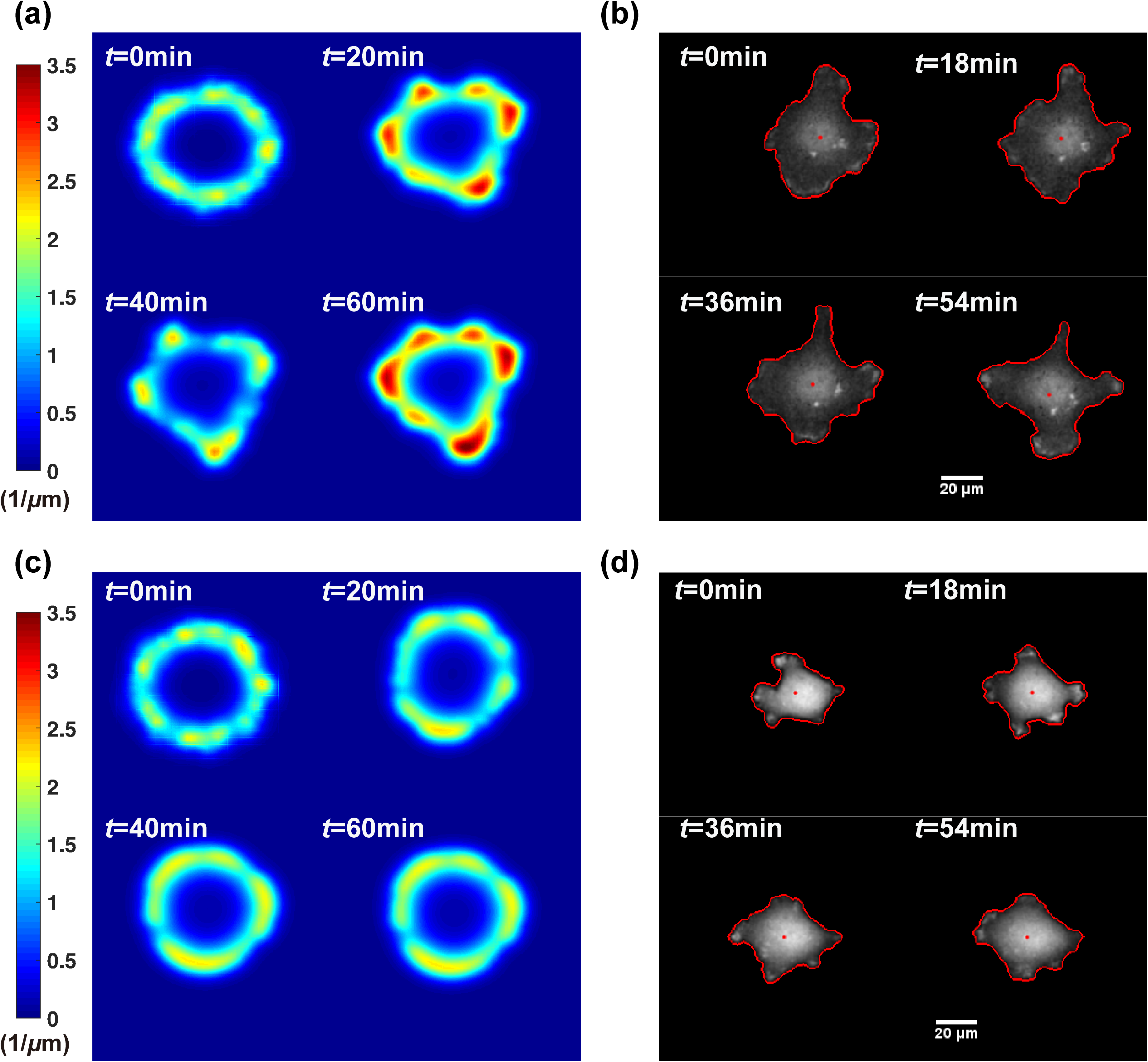
Comparison of cell deformation under random internal noise *in silico* and *in vitro*. **(a)** The cell membrane has evident deformation under low cell tension of *σ* = 0.005 *pN*/*μm*. Color-bar on the left represents concentration of F-actin. **(b)** MDA-MB-231 cells are more flexible in the hypertonic condition *in vitro*. The shape of cells is outlined in red with dots showing mass centroids of cells. Meanwhile, the brighter area indicates a higher concentration of PI3K which is an upstream kinase of F-actin. Scale bar: 20*μm*. **(c)** Cells lack dominant protrusions under a large tension of 0.015 *pN*/*μm*. Likewise, the colorbar shows the concentration of F-actin. **(d)** MDA-MB-231 cells are more rigid in hypo-condition *in vitro*. Cells are outlined in red with dots standing for the mass centroids of cells. Likewise, the brighter area indicates a higher concentration of PI3K. Scale bar: 20*μm*.

We confirm these predictions of our model by imaging on MDA-MB-231 cells in hypertonic and hypotonic conditions *in vitro*, i.e., low and high cell tension, respectively.

Both the results *in silico* and *in vitro* reveal that cell tension could be a shape regulator under the condition of random internal noise and from a qualitative point of view. Furthermore, our numerical results shed light on further explorations and bring a series of questions that require more detailed and quantitative researches, such as how to measure cell migration ability, what is the relationship between migration speed and cell tension and so on.

### Various migration modes

Cell migration displays diverse patterns, depending on the strength and spatiotemporal distribution of stimuli as well as cell tension. Under random stimuli, our cell migration model generates three cell migration modes. Notably, this is a ubiquitous phenomenon at different cell tension (initial line tension) by adjusting suitable protrusive coefficients.

The high migration ability pattern is the most common migrating behavior (8 out of 10 simulations at low tension of 0.2 *pN*/*μm*) induced by random internal noise (Fig. 3 a). Due to non-uniformity of noise, the conversion rate from Rac-GDP to Rac-GTP is accelerated in some areas of the cell while inhibited in others. The amplification of such an effect results in a polarized distribution of F-actin preceding the formation of protrusions. Cells then move along the major direction where the dominant protrusion exists. Such a dominant protrusion disappears and reappears in different directions. Hence the migration direction of cells varies randomly accordingly.

**Fig. 3.**
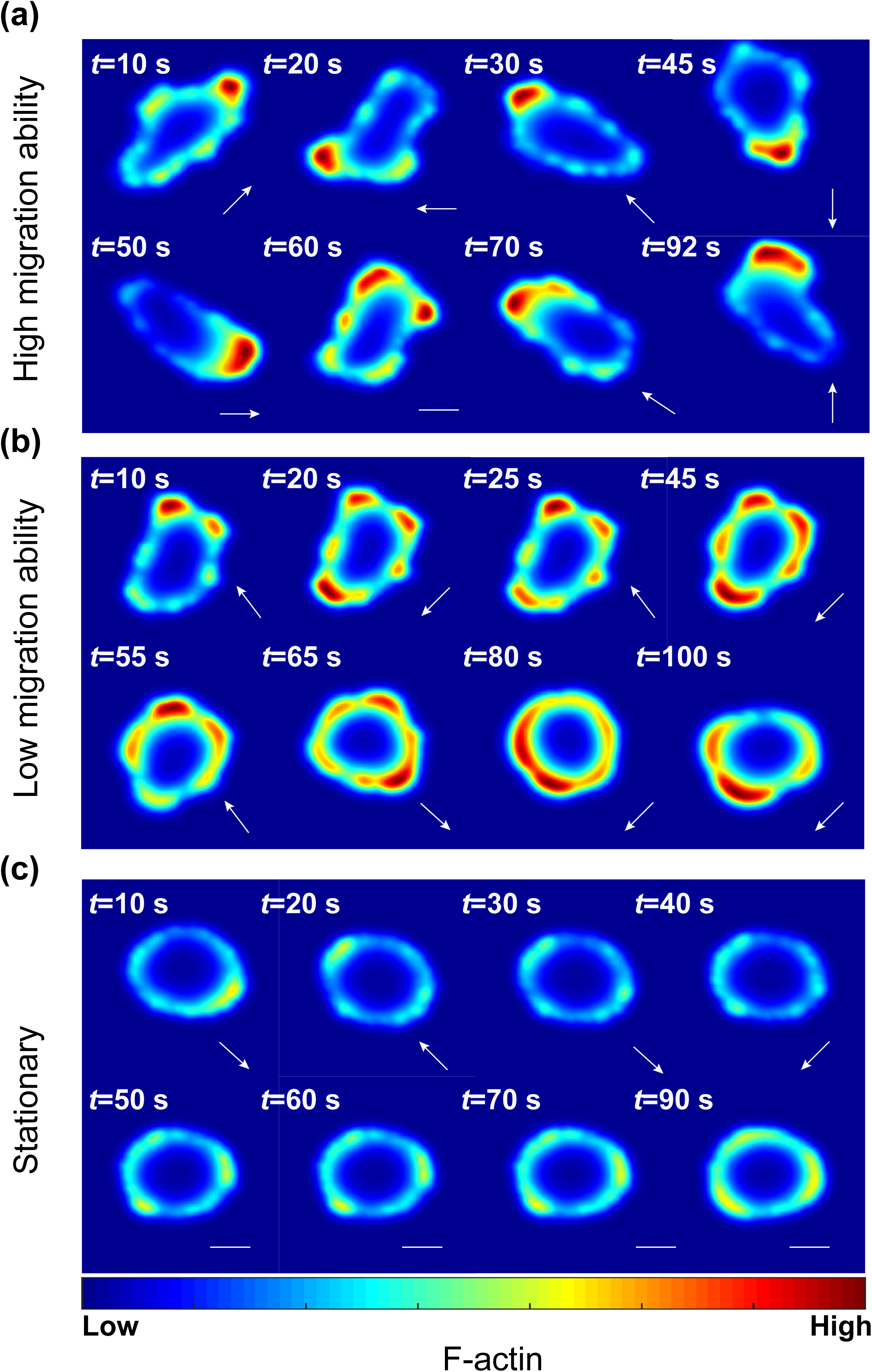
Three cell migration modes generated with the cell-migration model. **(a)** Representative time series of the high migration mode emerges due to the translocation of polarity patterns when *σ* is also chosen as 0.2 *pN*/*μm*. White bar stands for the tendency of staying at rest state. **(b)** Representative time series of the low migration ability mode as the parameter of initial line tension *σ* is attuned as 0.2 *pN*/*μm*. The white arrow at bottom right corner indicates the instant direction for cell movement. **(c)** Stationary migration mode presents when *σ* increases to 0.8 *pN*/*μm*. Cells tend to keep a regular morphology with a weak concentration of F-actin and either possess a stable movement or be still.

Cells also show low migration ability patterns with ~20% probability at low tension of 0.2 *pN*/*μm* in the simulations (Fig. 3 b). One interesting property of this pattern is that the polarized distribution of F-actin at opposite sides of the cell appears and vanishes periodically. Thus under the influence of oscillatory polarized distributions, cell migrates in a periodic pattern because the protrusive force is proportional to the concentration of F-actin. The oscillation of movement can be disrupted as cells are capable of re-polarizing and generating protrusions along other directions. When other protrusions are in a dominant position, the equilibrium of the oscillatory pattern is broke and the cell moves toward a new direction (Fig. 3 b).

Stationary patterns are generated when cell tension is 0.8 *pN*/*μm* (7 out of 10 simulations). Since high tension increases the threshold of polarization, compared to cells in low tension, the maximum concentration of F-actin is highly inhibited, although polarized distribution of F-actin could still occasionally show up in some parts of membrane due to non-homogeneity of random noise, and cell membrane do not show obvious deformations (Fig. 3 c).

These simulated patterns are consistent with the observed movements of spontaneous polarized MDA-MB-231 cells without any external gradient stimuli (Movie S3-S5).

### High cell tension inhibits movement under persistent external stimuli

Our models also show other migration patterns such as the persistent pattern given the presence of external stimuli (S2 Fig. c). As shown in Figs. 4 c-d, given external stimuli in a fixed direction, cells are stretched and move along the direction of external stimuli.

**Fig. 4.**
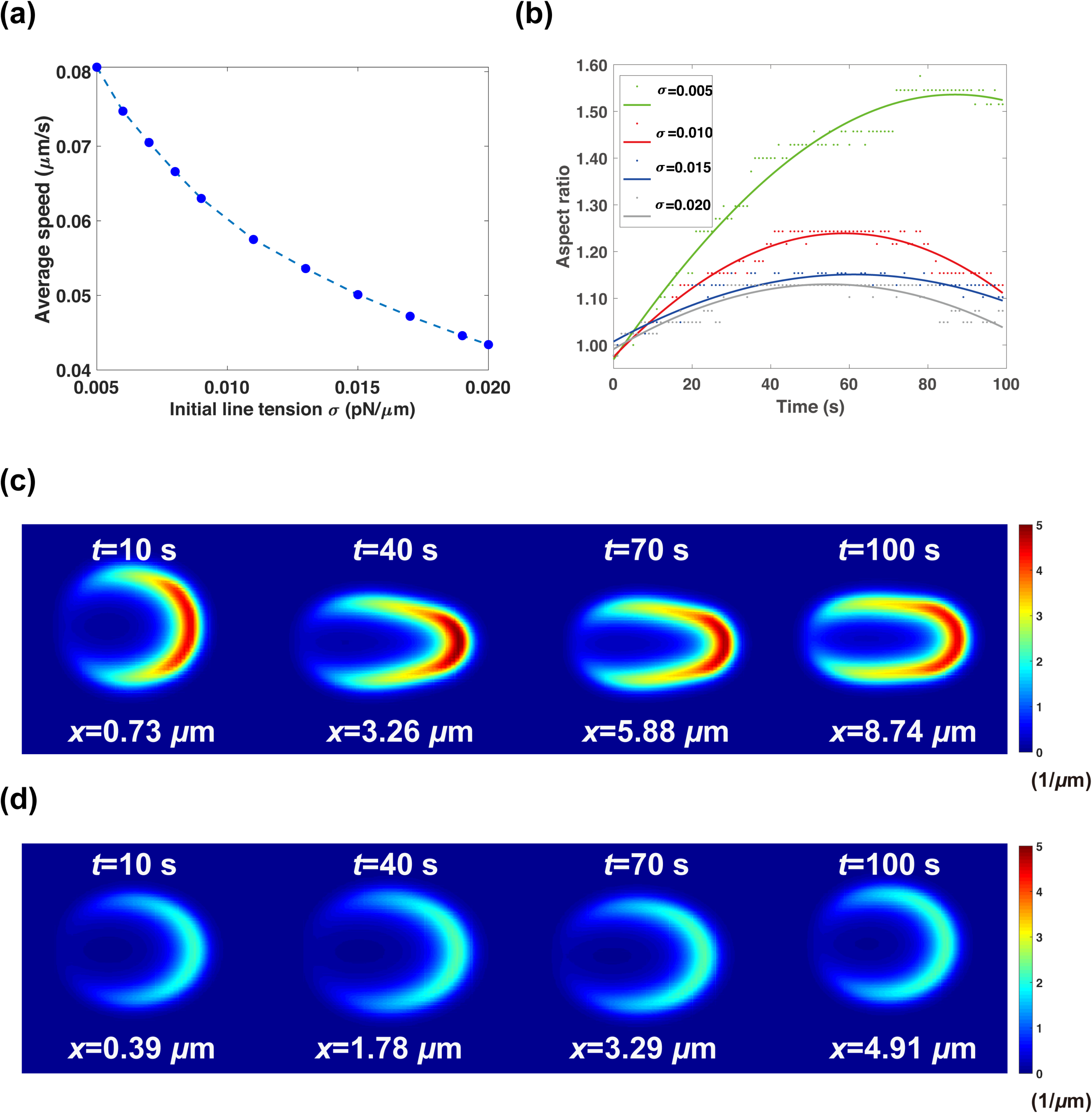
Cell migration ability under persistent external stimuli. **(a)** The average speed as a function of different initial line tension. The average speed is computed by the ratio between traveling distance and total simulation time. **(b)** Change of the aspect ratio of cells under four representative values of initial line tension. The aspect ratio measures the ratio between the maximum and minimum radius of cells is dependent on time during movement. Dots in green, red, blue and grey mean the sample points collected via simulations, while the corresponding smoothing sinusoidal curves are fitted with an average coefficient of determination *R*^2^ = 0.8855. **(c)** Cell migration ability and spatial profile of F-actin in low tension. *σ* is at the value of 0.005 *pN*/*μm*, while the color-bar indicates the concentration of F-actin. **(d)** Cell barely changes its shape with the maximum concentration of F-actin drops significantly at high cell tension of 0.02 *pN*/*μm*.

We find cell tension acts as an inhibitor for the movement as the average speed drops nearly a half from 0.08 *μm*/*s* at 0.005 *pN*/*μm* to 0.044 *μm*/*s* at 0.02 *pN*/*μm* (Fig. 4 a). This is because of the following reasons. First, cells tend to stay unpolarized under higher cell tension (Fig. 1 d). Second, the polarized cells lose propensity of changing their shape to be elongated under higher cell tension (Figs. 4 b-c). But based on our simulations, elongated cells with large aspect ratio move faster (S2 Fig. a). As shown in S2 Fig. a the average speed is linearly dependent on the aspect ratio with a goodness of fit as high as 0.95. This strong correlation is reasonable. As shown in the simulation, cells polarize in response to the external spatial cue, newly polymerized F-actin leads to rearrangement of cytoskeleton structure, cells elongate along the polarization direction and are poised for movement. Then they migrate by the collective efforts of protrusion and retraction (Movie S6-S7). Last, the persistent time of cell movement decreases under higher cell tension (S2 Fig. b). Polarized cells only maintain the shape with a high aspect ratio and keep moving in a certain duration, which is denoted as the persistent time. When cells undergo deformation and crawling, the total concentration of F-actin accumulates because of the positive feedback of Rac-GTP. Meantime, the cell perimeter notably increases as cell extends. These two factors coordinately elevate cell tension inhibiting the polymerization of F-actin. As a result, the maximum concentration of F-actin gradually drops and eventually redistributes evenly around the cell. And the cell finishes migration with the decreased aspect ratio. Since cells with lower cell tension are more inclined to deform in shape (Fig. 3), it takes a much longer time for cells with lower cell tension to return its shape to initial round disk (Fig. 4 b, *σ* = 0.005 *pN*/*μm*).

Our simulation result shows in the circumstance of persistent external stimuli, cell migration ability is restricted under high tension as movement is merely driven by a solo lamellipodium. This result is consistent with the published result showing elevated membrane tension confines actin assembly which results in larger leading edge and defective chemotaxis (37).

### Migration ability is maximized at intermediate cell tension under random internal noise

Unlike exposed to external stimuli with migration driven by protrusions directly toward the source of stimuli, cells are capable of generating multiple lamellipodiums (or filopodiums) spontaneously under the regulation of random internal noise, which is more frequently to be observed *in vitro*. Surprisingly, our cell migration model shows a non-monotonous relationship in terms of instant speed and cell tension within the range of [0.005 *pN*/*μm*, 0.015 *pN*/*μm*].

As shown in the simulation, when cell tension is low, cells are easy to polarize and more likely to generate multiple protrusions under small perturbations from internal noise. These lamellipodiums on random directions offset the momentum in the movement on principle direction. With the increase of cell tension, some protrusions which are induced by small perturbations are inhibited as tension stabilizes polarity and therefore cells are better streamlined. Cells with a streamlined body reach a higher migration ability. Thus cells acquire fast instant speed. However, if cell tension continues to increase, the stiffness of membrane prohibits F-actin polymerization (Fig. 5 a). In our simulations, we take the mean value of instant speed during the whole simulation period and also average on all samples. A similar trend is obtained with a measurement of average speed when initial line tension is within the range of [0.2 *pN*/*μm*, 0.8 *pN*/*μm*] (MCF-7 cells). The optimized average speed arises around *σ* = 0.4 *pN*/*μm* (Fig. 5b).

**Fig. 5.**
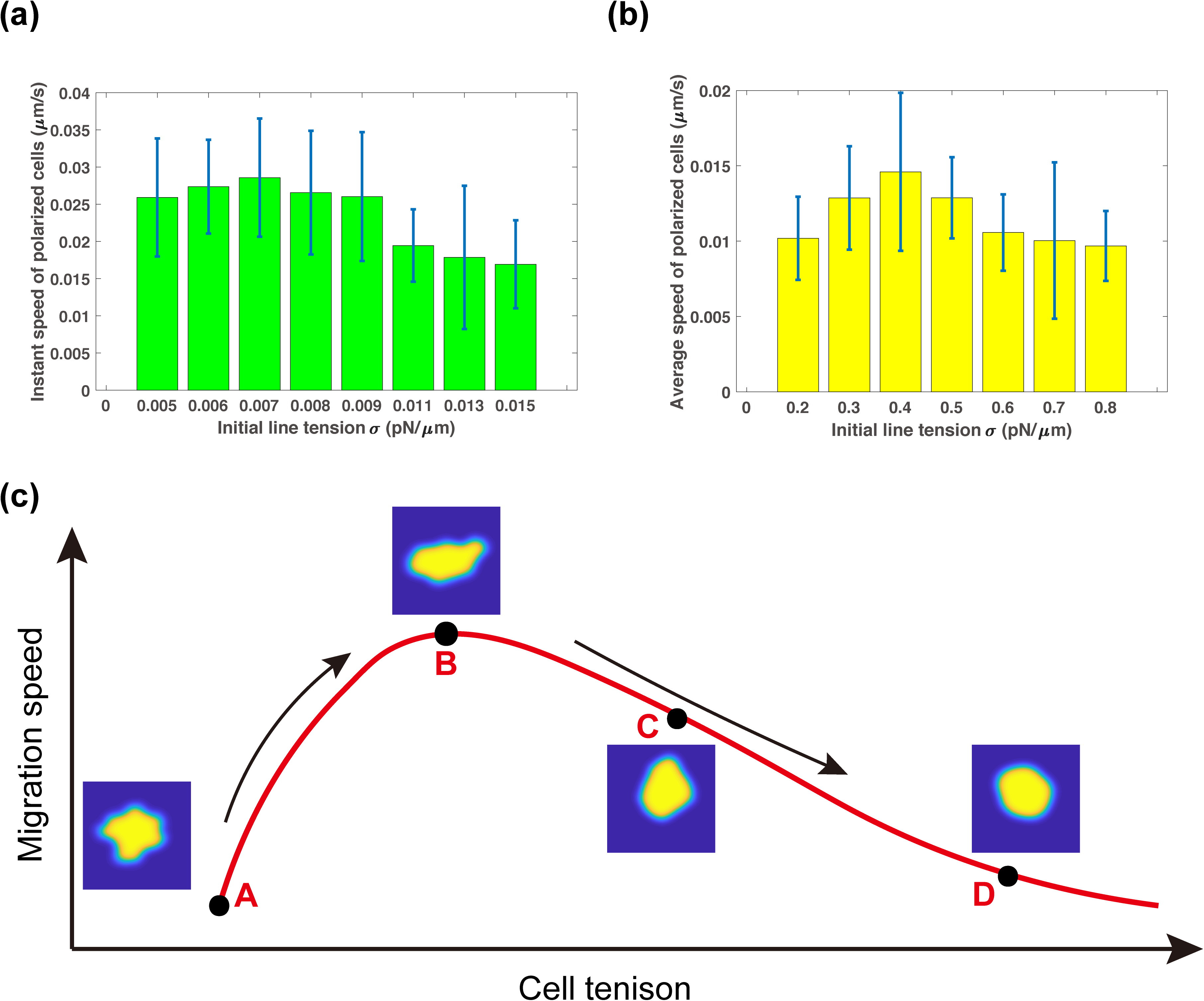
Non-monotonous relationship between the migration speed and cell tension under random internal noise. **(a)** The instant speed of polarized cells as a function of initial line tension *σ* within the range of [0.005 *pN*/*μm*, 0.015 *pN*/*μm*]. The instant speed is calculated as the difference of the centroids of cells at each time step. Error bars represent the standard deviations of the instant speed under sample size *N* = 20. **(b)** The average speed of polarized cells as a function of initial line tension *σ* within the range of [0.2 *pN*/*μm*, 0.8 *pN*/*μm*]. Since only a fraction of cells are polarized (Fig. 1d), the data for the average speed is recorded only from those polarized cells. Error bars represent the standard deviation. **(c)** A hypothetical curve regarding migration speed and cell tension. The black arrows indicate the tendency of migration speed. Points A to E are five representative phases under tension regulation. Point A is the initial cell shape while B to E stand for different cell morphologies under different cell tension.

We summarize the bell curve regarding cell migration speed and tension from the simulation (Fig. 5 c). These simulated results are consistent with previously published experimental results. Batchelder *et. al* used a simple model of *C. elegans* sperm cell to quantify the relation between membrane tension and cell migration speed. They surprisingly find that membrane tension reduction is correlated with a decrease in cell displacement speed, whereas an increase in membrane tension enhances motility (36). Similar results are observed in human neutrophils (5). Likewise, cell movement is impeded by multiple lamellipodia in moving keratocytes (6) and protruding fibroblasts (38). From the observations of both cases, cells are more streamlined under a relatively high cell tension. It is hypothesized that membrane tension can drive coalescence of protrusions and thus enhance motility and a non-monotone curve between tension and migration speed is also expected (14, 36). More systematic tests on the simulation results remained to be performed in future studies.

## Discussion

Cell migration is orchestrated by complicated mechanochemical systems. It entails F-actin polymerization and morphological change via combined effects from various forces, such as protrusive force, retraction, cell tension and adhesion. Here protrusive force and retraction are connected with activities of polarity marker proteins Rac-GTPase and protein motor myosin respectively. Cell tension consists of cortical tension and membrane tension. In response to external stimuli or/and random internal noise, polarity proteins accumulate on certain cell edges under positive feedbacks to form a polarized distribution. The cytoskeleton is rearranged to deform cell shape under the inhibition from cell tension. Cells move with the synergy of protrusions such as lamellipodia and retraction. Different migration modes emerge depending on the cell type, extracellular matrix, and stimuli. It is difficult to model a variety of phenomena of cell migration. Our models focus on the coupling between cell tension and biochemical factors including Rac-GTPase, F-actin, and myosin. In our work, tension is proportional to the total concentration of polymerized F-actin. While on the other hand, it is also used as a long-range inhibitor to deactivate F-actin activities. We describe mechanical properties via phase field model and use reaction-diffusion equations to express biochemical systems. Our models successfully reveal how cell tension affects cell polarization and movement. In simulations, our models recapitulate common features of polarization (Fig. 1). Based on our models, cells utilize one particular parameter initial line tension, in addition to the chemical systems, to regulate cell morphological change through competitions of protrusions (Fig. 2).

Cell tension is determined by both the stiffness of membrane and polymerization of actin filaments. The rigidity of the membrane is dependent on molecular arrangement and it varies from cell to cell. In our model, that the stiffness of membrane can be altered by attuning initial line tension σ. Different values of σ can represent various cell types, e.g., we select σ = 0.2 *pN*/*μm* and σ = 0.8 *pN*/*μm* to stand for cancer stem cells and non-stem cancer cells in MCF-7 cell line. Meanwhile, σ = 0.005 *pN*/*μm* is selected to represent MDA-MB-231 cells in hypertonic condition while σ = 0.015 *pN*/*μm* for those cells in hypotonic condition. Notably, after taking into the cortical tension into consideration, within the range of [0.005 *pN*/*μm*, 0.015 *pN*/*μm*], the effective cell tension σ′ is around 10 *pN*/*μm* to 25 *pN*/*μm* and σ′ soars to around 70 *pN*/*μm* to 140 *pN*/*μm* when parameter of initial line tension is taken as [0.2 *pN*/*μm*, 0.8 *pN*/*μm*] based on calculation of *mt*(*f, ϕ*) in Eq. 8, which are consistent with the measurement results on the other cell types (14). To ensure membrane deformation under different initial line tension σ, we could adjust one parameter, the protrusive coefficient. As the cell membrane deformability depends on the strength of protrusive force, which is proportional to the concentration of Factin but the synthesis F-actin is inhibited by cell tension.

Our model generates representative migration modes. When there is only random internal noise, a highly motile mode is the most common at low cell tension. Whereas cells prefer to be stationary at high tension (Fig. 3). Given external stimuli without random internal noise, cells show a persistent motion. These results are consistent with the experimental results. We further investigate how tension affects cell movement. We confirm that at cell tension inhibits the formation of protrusion, hence the cell with only one protrusion under a strong external stimulus show reduced migration ability as cell tension is high (Fig. 4 a, c-d, SI Fig. 2 a). Interestingly, when internal random noise dominates and cells form multiple protrusions, we discover a biphasic non-monotonous effect of tension regulation on cell migration ability, i.e., cells achieve the maximum migration ability at intermediate tension. This result is consistent with a previously proposed hypothesis (14).

Our proposed model provides a minimal system that cell tension regulates migration ability. For simplification, the current model neglects influence from the extracellular matrix (ECM) to movement. ECM activates signal transduction in cells and when cell membrane deforms, the contacting area between membrane and substrate is also changing. Besides, increasing contacts between cells and the extracellular matrix could increase the stiffness and viscosity of the actin cytoskeleton (39). Hence a more comprehensive model can be constructed to describe how ECM, mechanical and biochemical signals interact with each other. With this mode, more migration modes such as directed cell migration under topographic ECM gradients (40) and oscillation patterns (41) could be investigated. Meanwhile, we only focus on instant and average speed to measure migration ability in our model. It will be interesting to use other characteristics such as cell turning to measure migration ability in the future.

## Author Contribution

Conceptualization: KT FL LZ WW

Investigation: KT JW XK

Methodology: KT JW FL LZ

Supervision: FL LZ

Writing - original draft: FL LZ KT JW

## Acknowledgments

We thank Yihan Lin lab for providing the transfected cell lines used in this study. This work was supported by the National Natural Science Foundation of China (11622102, 11861130351,11434001,31670852).

## Supporting Materials

Supporting Materials include two supplementary Figures, seven supplementary Movies, and one supplementary Table. References (42, 43) are also cited in the Supporting Materials.

